# Validation of GCN5L1/BLOC1S1/BLOS1 Antibodies Using Knockout Cells and Tissue

**DOI:** 10.1101/2023.07.21.550091

**Authors:** Paramesha Bugga, Michael W. Stoner, Janet R. Manning, Bellina A.S. Mushala, Dharendra Thapa, Iain Scott

**Affiliations:** Vascular Medicine Institute, University of Pittsburgh, Pittsburgh, PA 15261; Center for Metabolism and Mitochondrial Medicine, University of Pittsburgh, Pittsburgh, PA 15261; Division of Cardiology, Department of Medicine, University of Pittsburgh, Pittsburgh, PA 15261; Department of Human Performance - Exercise Physiology, West Virginia University, Morgantown, WV 26506

**Keywords:** GCN5L1, BLOC1S1, BLOS1, Antibody, Mitochondria, Autophagy, Lysosome

## Abstract

GCN5L1, also known as BLOC1S1 and BLOS1, is a small intracellular protein involved in a number of key biological processes. Over the last decade, GCN5L1 has been implicated in the regulation of protein lysine acetylation, energy metabolism, endo-lysosomal function, and cellular immune pathways. An increasing number of published papers have used commercially-available reagents to interrogate GCN5L1 function. However, in many cases these reagents have not been rigorously validated, leading to potentially misleading results. In this report we tested several commercially-available antibodies for GCN5L1, and found that two-thirds of those available did not unambiguously detect the protein by western blot in cultured mouse cells or *ex vivo* liver tissue. These data suggest that previously published studies which used these unverified antibodies to measure GCN5L1 protein abundance, in the absence of other independent methods of corroboration, should be interpreted with appropriate caution.

## Introduction

GCN5L1 (also known as BLOC1S1 and BLOS1; hereafter referred to as GCN5L1) is a highly conserved small protein of 15-17 kD, with homologues found throughout the plant and animal kingdom (Scott et al, 2018). First identified as a human cDNA with partial sequence homology to the lysine acetyltransferase GCN5/KAT2A (Inoue et al, 1996), GCN5L1 has been implicated in the regulation of a number of key cellular processes. Based on its sequence homology to bacterial and human acetyltransferases, and its cellular localization to mitochondria, we identified a role for GCN5L1 in the lysine acetylation of mitochondrial proteins in the electron transport chain, the oxidation of fuel substrates, and mitochondrial DNA replication/maintenance (Scott et al, 2012; Thapa et al, 2017; Thapa et al, 2018; Thapa et al, 2022; Manning et al, 2022; Zhang et al, 2023). Subsequently, other groups have confirmed this function for GCN5L1, demonstrating that its loss impacts mitochondrial glutaminase activity, reactive oxygen species detoxification, and TFAM acetylation (Lv et al, 2019; Lv et al, 2021; Zhang et al, 2022; Lv et al, 2022).

Another major strand of GCN5L1 research has focused on its role in endo-lysosomal regulation. GCN5L1 was identified in 2004 as a component of the BLOC1 complex, which regulates the biogenesis of lysosome-related organelles (Starcevic and Dell’Angelica, 2004). Subsequent studies have shown that GCN5L1 is involved in endosome-to-vacuole transport in Arabidopsis (Cui et al, 2010), the development of lysosome-related organelles in Zebrafish (Chen et al, 2018), and lysosomal trafficking in mouse liver (Wu et al, 2018). More recently, these endo-lysosomal processes have implicated GCN5L1 in the control of stress-related lysosomal positioning (Bae et al, 2019), LDL receptor recycling (Zhang et al, 2020), bacterial innate immune pathways (Wells et al, 2022), and hepatic lysosomal lipolysis (Wu et al, 2023).

The increased interest in GCN5L1 over the last decade has led to an explosion in the number of published studies, many of which rely on the use of commercial reagents to interrogate GCN5L1 biology. In terms of protein abundance, several commercial antibodies are available for GCN5L1 for use in western blotting. However, the majority of these commercial antibodies have not undergone rigorous testing to determine their specificity and selectivity for GCN5L1. In this study, we attempted to validate commercial antibodies for GCN5L1 from Proteintech, Sigma Aldrich, and Santa Cruz Biotechnology, using qPCR-validated wildtype and GCN5L1 knockout cells and mouse tissue. Our results show that the majority of antibodies commercially available for GCN5L1, including those used in multiple published studies, are not specific for this protein.

## Materials and Methods

### Animal Care and Use

Male liver-specific GCN5L1 knockout (KO) and wildtype (WT) controls were bred as previously described (Wang et al, 2017). Animals were housed in the University of Pittsburgh animal facility under standard conditions with *ad libitum* access to water and food, and maintained on a constant 12-hour light/12-hour dark cycle. Mice were fed a control chow diet from weaning to 8-10 weeks of age, then euthanized by CO_2_ asphyxiation/cervical dislocation. Experiments were conducted in compliance with National Institutes of Health guidelines, and followed procedures approved by the University of Pittsburgh Institutional Animal Care and Use Committee.

### Cell Culture Conditions, Protein Isolation, and Immunoblotting

Wildtype and GCN5L1 knockout mouse embryonic fibroblast (MEF) cells were cultured at 37 °C/5% CO_2_ in DMEM with antibiotic/antimycotic as previously described (Scott et al, 2014). Cells were washed in ice-cold 1X PBS and harvested by scraping using a cell lifter. Cells were spun at 500 *g* for 5 min, and the pellets lysed on ice in 1X CHAPS buffer for 1 hour (1% CHAPS, 150 mM NaCl, 10 mM HEPES, pH 7.4). Homogenates were spun at 10,000 *g* at 4°C for 10 min, and the supernatants collected for immunoblotting.

Liver tissues were rapidly harvested following euthanasia, weighed and flash-frozen in liquid nitrogen. Tissues were minced and lysed in 1X CHAPS buffer (1% CHAPS, 150 mM NaCl, 10 mM HEPES, pH 7.4) using a VWR 4-Place Mini Bead Mill, then incubated on ice for ∼2.5 hours. Homogenates were spun at 10,000 *g* at 4°C for 10 min, and the supernatants collected for immunoblotting.

For immunoblotting, protein lysates were quantitated using a BioDrop μLITE Analyzer, prepared in LDS sample buffer, separated using Bolt SDS-PAGE 12% Bis-Tris Plus gels, and transferred to nitrocellulose membranes (all Invitrogen). Membranes were blocked using Intercept Blocking Buffer and incubated overnight in the following primary antibodies: rabbit GAPDH (2118S) from Cell Signaling Technology; rabbit BLOC1S1 (19687-1-AP) from Proteintech; mouse BLOS1 (sc-515444) from Santa Cruz Biotechnology; rabbit BLOC1S1 antibody (HPA021381) from Sigma Aldrich; and custom-generated GCN5L1 (Covance) as previously reported (Scott et al, 2012). Protein loading was confirmed using GAPDH as a loading control. Fluorescent anti-goat or anti-rabbit secondary antibodies (red, 700 nm; green, 800 nm) from LiCor were used to detect expression levels. Images were obtained using Licor Odyssey CLx System and protein densitometry was measured using the LiCor Image Studio Lite Ver. 5.2 Software.

### RNA Isolation and Quantitative RT-PCR

Total RNA was isolated from liver tissue and MEF cells using the RNeasy Mini Kit (Qiagen). RNA was quantitated, and 300-1000 ng was used to generate cDNA using Maxima Reverse Transcriptase Kit (ThermoFisher). Quantitative real-time PCR was performed using 1X Power SYBR-Green PCR Master Mix (ThermoFisher), with validated gene-specific primers (Qiagen) on the QuantStudio™ 5 System. Primers used were QT01063069 (*Gcn5l1/Bloc1s1*) and QT01658692 (*Gapdh)*. Each experiment was performed in triplicate and data were analyzed via the ΔC_t_ method using *Gapdh* as the reference gene.

### Statistics

Means ± SEM were calculated for all data sets. Comparisons between single variable groups were made using unpaired, two-tailed Student’s T-Tests. *P* ≤ 0.05 was considered statistically significant. Statistical analyses were performed using GraphPad Prism 9.0 Software.

## Results

### Detection of GCN5L1 protein abundance in mouse embryonic fibroblasts

Mouse embryonic fibroblasts (MEFs) lacking GCN5L1 expression display decreased mitochondrial protein acetylation levels and increased mitochondrial protein turnover rates (Webster et al, 2013; Scott et al, 2014). We used these cells to test whether different custom and commercial antibodies could unambiguously detect GCN5L1 proteins levels. Firstly, we validated gene expression in wildtype (WT) and GCN5L1 knockout (KO) MEF cells using qPCR, and confirmed that *Gcn5l1* transcript levels were essentially undetectable in KO MEFs (**Figure 1**). We then examined GCN5L1 protein abundance by western blot, using either a custom polyclonal GCN5L1 antibody (generated by Covance; Scott et al, 2012), or commercial polyclonal antibodies obtained from Proteintech, Sigma Aldrich, and Santa Cruz Biotechnology. The peptide immunogens used to generate each of the GCN5L1 antibodies (other than the Proteintech antibody, which was not reported by the manufacturer) are shown in **Figure 2**. In agreement with previous reports (Scott et al, 2012; Scott et al, 2014), there was effectively no immunoreactivity at the correct molecular weight (∼15 kD) in GCN5L1 KO MEFs using the custom Covance antibody (**Figure 3A**). The same result was obtained using the Sigma Aldrich commercial antibody, which showed immunoreactivity at the correct molecular weight in only WT MEF cells (**Figure 3C**).

**Figure 1.**
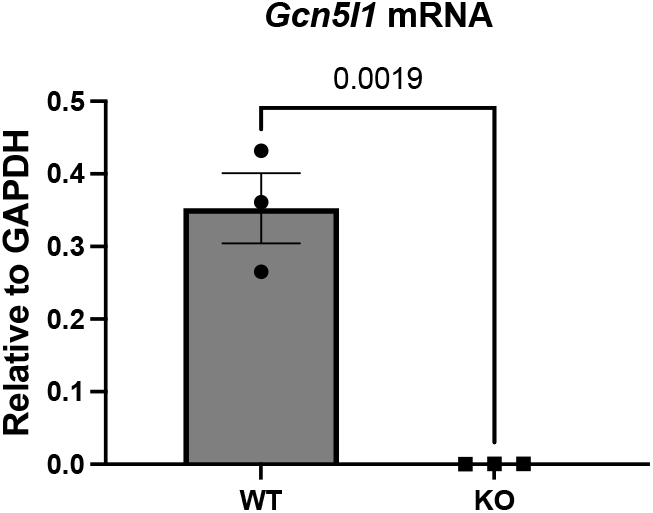
GCN5L1 transcript levels in wildtype (WT) and knockout (KO) mouse embryonic fibroblasts (MEFs). Quantitative PCR (qPCR) was used to confirm that GCN5L1 gene expression was absent in KO MEF cells. N = 3.

**Figure 2.**
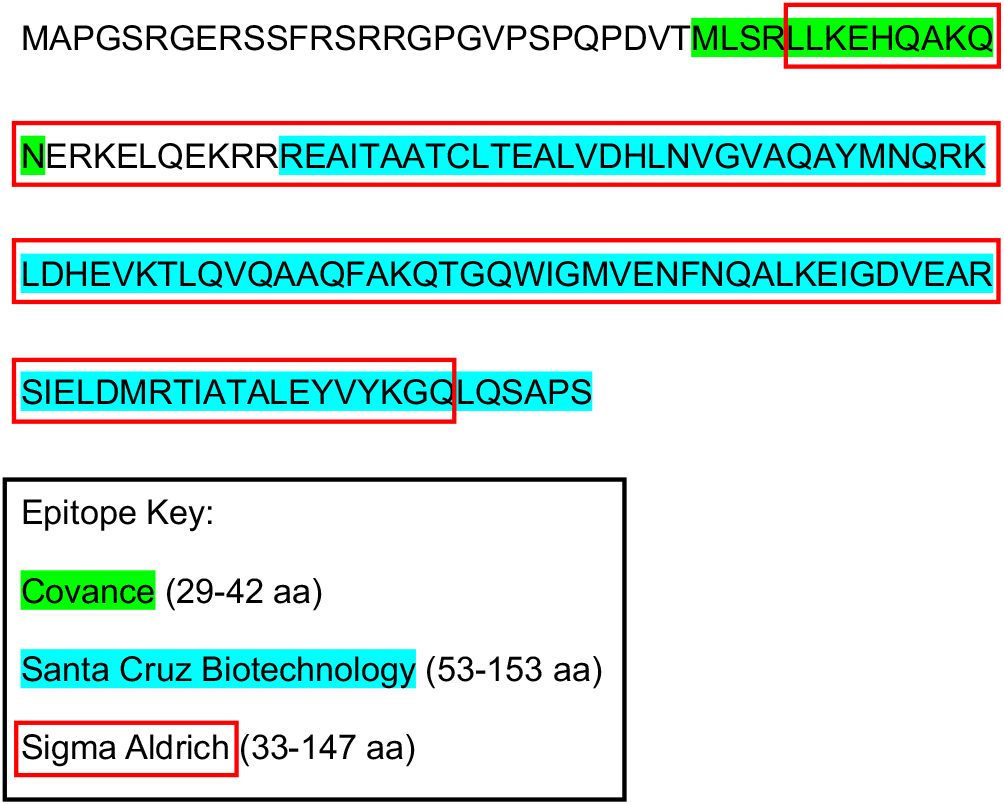
Peptide immunogens used to generate GCN5L1 antibodies. The epitopes of each of the antibodies used in this study mapped to human GCN5L1 (1-153 aa). The peptide immunogen sequence for the Proteintech antibody was not reported.

**Figure 3.**
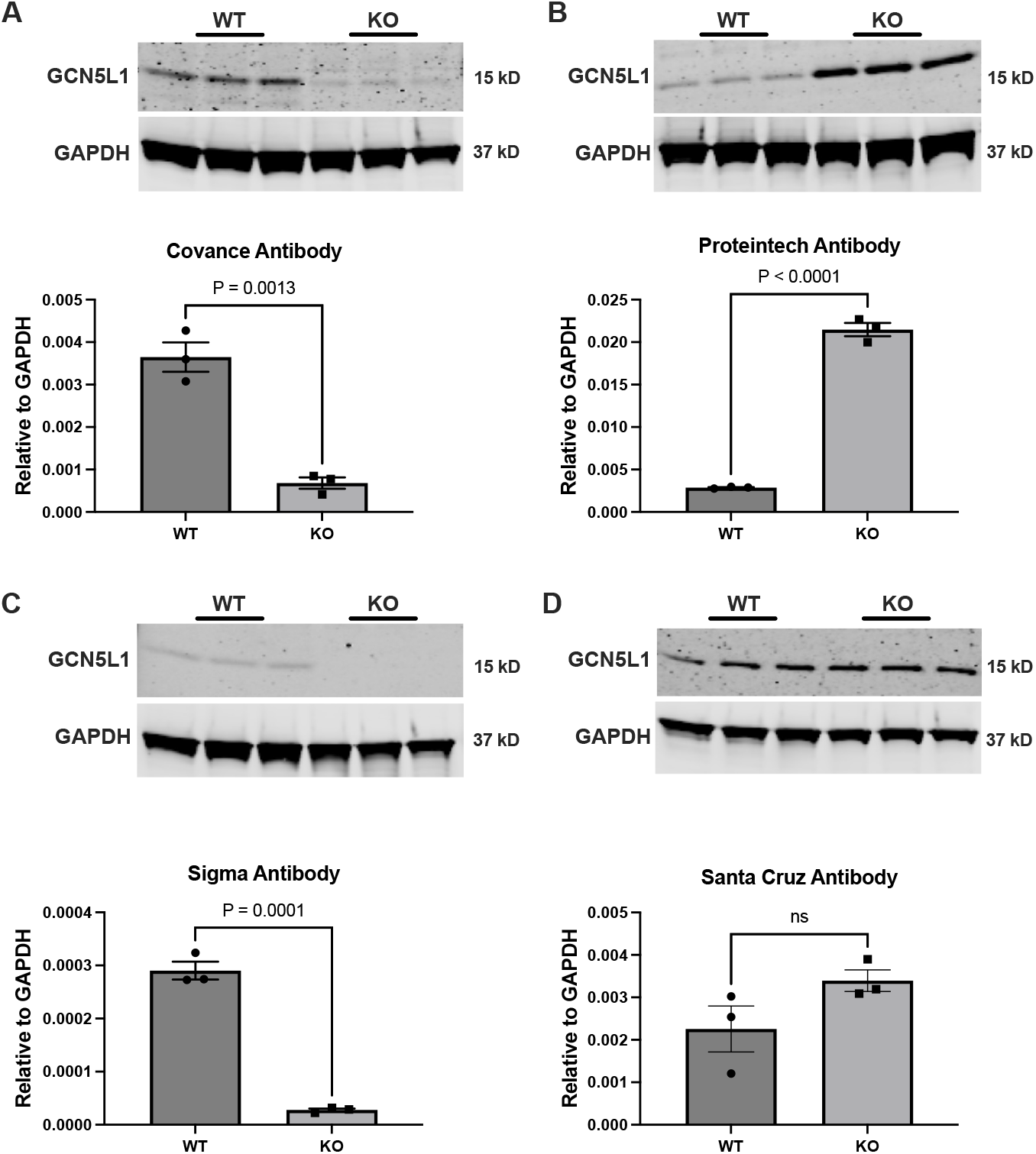
Measurement of GCN5L1 protein abundance in wildtype (WT) and knockout (KO) mouse embryonic fibroblasts (MEFs). GCN5L1 protein was essentially undetectable in KO MEFs using the Covance and Sigma antibodies (A,C). In contrast, an immunoreactive band at the correct molecular weight for GCN5L1 was detected in validated KO MEFs using both the Proteintech and Santa Cruz antibodies (B,D). N = 3.

In contrast to these expected results, we found that both the Proteintech and Santa Cruz Biotechnology commercial antibodies displayed robust immunoreactivity at the correct ∼15 kD molecular weight in both WT and KO MEFs (**Figure 3B,D**). This result was particularly striking using the Proteintech antibody, where KO cells actually showed a significant increase in abundance of the protein band at 15 kD (**Figure 3B**). From these results, we conclude that both the Proteintech and Santa Cruz Biotechnology commercial antibodies are unable to unambiguously detect GCN5L1 in MEF cells.

### Detection of GCN5L1 protein abundance in mouse liver tissue

Mouse livers with hepatocyte-specific deletion of GCN5L1 display reduced hepatic gluconeogenesis, aberrant lysosomal positioning, and increased fatty acid oxidation rates (Wang et al, 2017; Wu et al, 2018; Thapa et al, 2018). To validate our findings from MEF cells, we used liver tissue from WT and hepatocyte-specific GCN5L1 KO mice. In agreement with the MEF cells, there was almost no GCN5L1 mRNA detected in liver tissue from KO mice (**Figure 4**), with the residual expression likely to be from other liver cell types (Kupffer cells, hepatic stellate cells, etc.). We used the same antibodies to examine GCN5L1 protein abundance, and found good agreement with our results using MEF cells. The custom Covance and commercial Sigma Aldrich antibodies both detected a band of the correct molecular weight in WT tissue, which was absent in GCN5L1 KO livers (**Figure 5A,C**).

**Figure 4.**
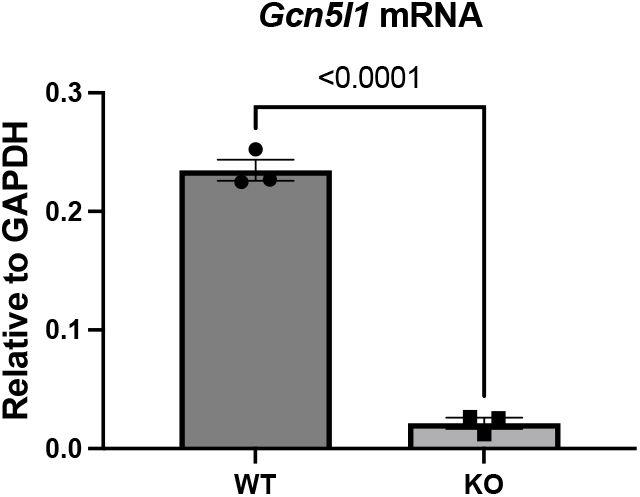
GCN5L1 transcript levels in wildtype (WT) and knockout (KO) mouse liver tissue. Quantitative PCR (qPCR) was used to confirm that GCN5L1 gene expression was absent in KO liver tissue. N = 3

**Figure 5.**
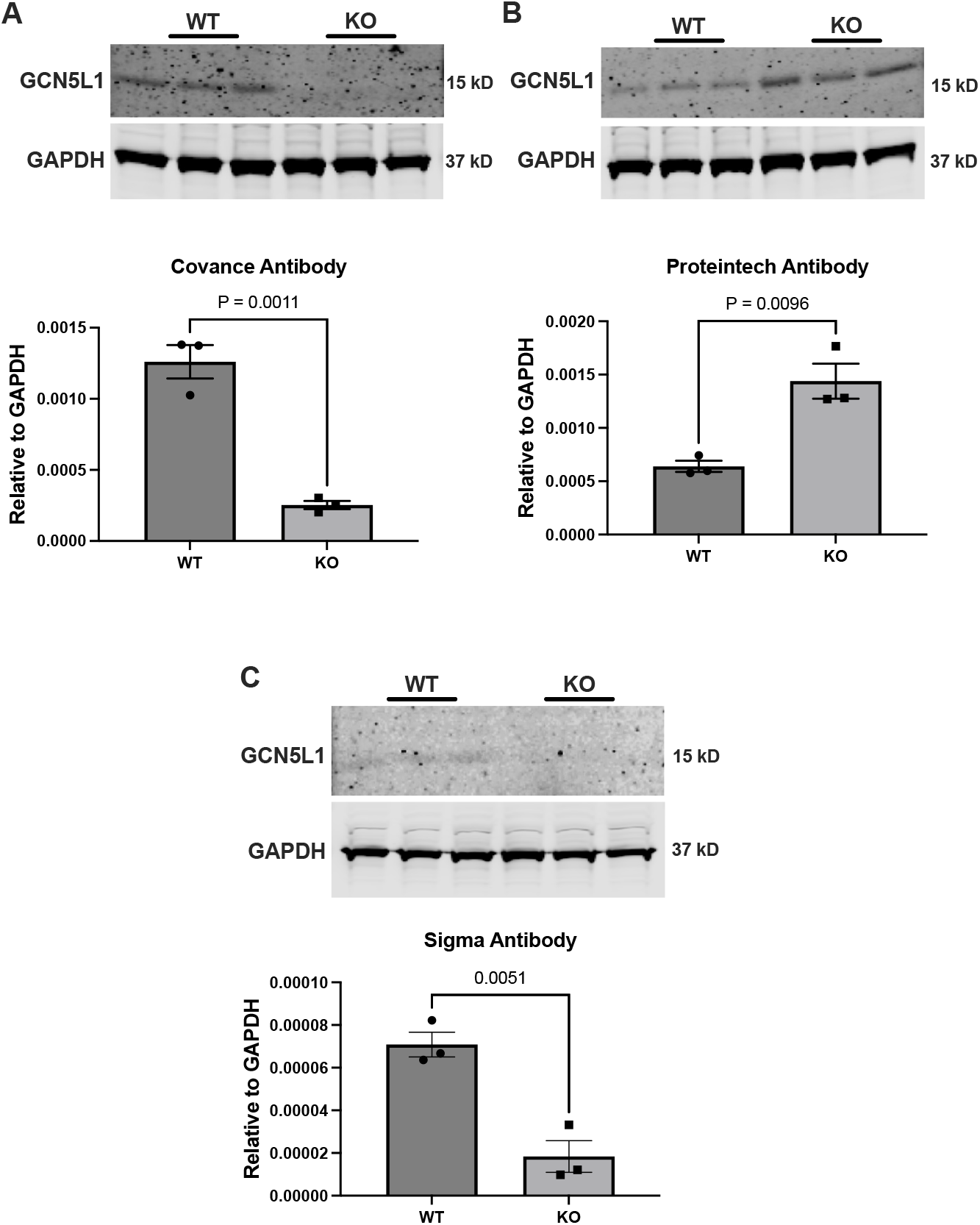
Measurement of GCN5L1 protein abundance in wildtype (WT) and knockout (KO) liver tissue. GCN5L1 protein was essentially undetectable in KO livers using the Covance and Sigma antibodies (A,C). In contrast, an immunoreactive band at the correct molecular weight for GCN5L1 was detected in validated KO livers using the Proteintech antibody (B). N = 3.

As with the MEF cells, the commercially-available Proteintech antibody gave the opposite result, detecting a band at the correct GCN5L1 molecular weight in both WT and KO livers (**Figure 5B**). As before, the Proteintech antibody appeared to detect a significantly-increased abundance of GCN5L1 protein in KO livers, despite an absence of GCN5L1 message detected by qPCR (**Figure 4**,**5**). Unfortunately, the Santa Cruz Biotechnology antibody was unable to detect GCN5L1 protein in either WT or KO liver samples, which prevented our analysis of its efficacy from *ex vivo* liver tissues (**Supplemental Figure 1**). From these data, we conclude that the GCN5L1 antibodies from Proteintech or Santa Cruz Biotechnology are not suitable for use in studies measuring GCN5L1 protein abundance.

## Discussion

Recent studies on GCN5L1 have determined that it plays a key role in a number of important cellular processes. To allow further rigorous studies on GCN5L1 function to be performed, it is crucial that researchers in the field have access to validated reagents that can unambiguously measure features of GCN5L1 biology. To this end, we examined the specificity of the three most frequently used commercial antibodies for GCN5L1, and tested them against a previously validated custom antibody in WT and KO MEF cells and liver tissue (Scott et al, 2012; Scott et al, 2014; Wang et al, 2017; Wu et al, 2018; Thapa et al, 2018).

Our results show that the GCN5L1 antibody from Sigma Aldrich performed as well as our previously-validated custom antibody made by Covance (Scott et al, 2012). This suggests that the overlapping region of the two peptide immunogens for these antibodies (LLKEHQAKQN, 33-42 aa in human GCN5L1; **Figure 3**) is necessary for the development of a successful and specific GCN5L1 antibody. In contrast, the GCN5L1 antibodies produced by Proteintech and Santa Cruz Biotechnology were unable to unambiguously detect GCN5L1 in MEF cells or mouse liver tissue. Our results therefore suggest that two-thirds of the GCN5L1 antibodies available commercially are not specific for this protein. These findings highlight the need for rigorous testing of commercially-obtained reagents in GCN5L1-focused research, prior to their use in published studies. Furthermore, these data suggest that a note of caution should be applied when interpreting any published studies that use these unvalidated antibodies to measure GCN5L1 protein levels, in the absence of another independent means of analysis.

Several proposals have been made to generate a consensus antibody validation strategy (see, e.g., Bordeaux et al, 2010; Uhlen et al, 2016). The latter proposal suggests five “pillars” of antibody validation, the first being whether genetic disruption (via genome editing or RNA interference) leads to the elimination or significant reduction of protein abundance by western blot. In the absence of this, another approach should be added, including the use of mass spectrometry, or the use of over-expressed epitope-tagged constructs to correlate with the endogenous protein levels, molecular weight, etc. (Uhlen et al, 2016).

In this study, we tested the four antibodies using qPCR-validated WT and GCN5L1 KO MEF cells and liver tissue. Several published studies on GCN5L1 have used the non-specific Proteintech or Santa Cruz Biotechnology antibodies in concert with siRNA knockdown to modulate GCN5L1 abundance. However, our results here suggest that this method did not sufficiently increase the rigor of these findings, as a protein band at the correct molecular weight was detected by these antibodies in validated GCN5L1-null samples. Therefore, we suggest that in addition to using genetic disruption methods to validate GCN5L1 antibodies, researchers in future studies should also include data to show that the genetic disruption itself has been successful, using other means, prior to publication.

## Author Contributions

Performed experiments: PB, MWS, JRM, BASM, DT. Designed experiments: PB, MWS, IS. Analyzed data: MWS, IS. Produced figures: MWS, IS. Wrote/edited manuscript: PB, DT, IS.

**Supplemental Figure 1.**
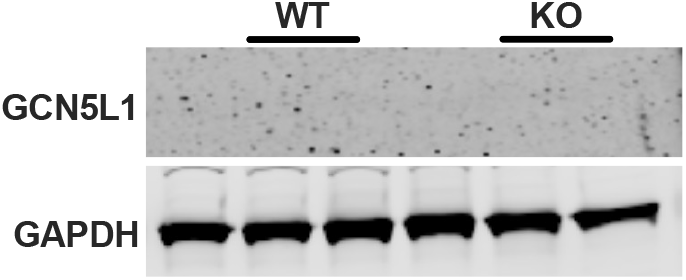
Santa Cruz Biotechnology antibody in liver tissue. GCN5L1 protein was not detectable in either wildtype (WT) or knockout (KO) liver tissue using the Santa Cruz antibody. N = 3.

